# Comparative genomics of Chinese and international isolates of *Escherichia albertii*: population structure and evolution of virulence and antimicrobial resistance

**DOI:** 10.1101/2021.02.01.429068

**Authors:** Lijuan Luo, Hong Wang, Michael Payne, Chelsea Liang, Li Bai, Han Zheng, Zhengdong Zhang, Ling Zhang, Xiaomei Zhang, Guodong yan, Nianli Zou, Xi Chen, Ziting Wan, Yanwen Xiong, Ruiting Lan, Qun Li

**Affiliations:** School of Biotechnology and Biomolecular Sciences, University of New South Wales, Sydney, New South Wales, Australia; Zigong Center for Disease Control and Prevention, Zigong, China; Division I of Risk Assessment, National Health Commission Key Laboratory of Food Safety Risk Assessment, Food Safety Research Unit (2019RU014) of Chinese Academy of Medical Science, China National Center for Food Safety Risk Assessment, Beijing, China; State Key Laboratory of Infectious Disease Prevention and Control, Collaborative Innovation Center for Diagnosis and Treatment of Infectious Diseases, National Institute for Communicable Disease Control and Prevention, Chinese Center for Disease Control and Prevention, Beijing, China

**Keywords:** *Escherichia albertii*, population structure, virulence, MDR, plasmid, prophage

## Abstract

*Escherichia albertii* is a newly recognized species in the genus *Escherichia* that causes diarrhea. The population structure, genetic diversity and genomic features has not been fully examined. Here, 169 *E. albertii* isolates from different sources and regions in China were sequenced and combined with 312 publicly available genomes for phylogenetic and genomic analyses. The *E. albertii* population was divided into 2 clades and 8 lineages, with lineage 3 (L3), L5 and L8 more common in China. Clinical isolates were observed in all clades/lineages. Virulence genes were found to be distributed differently among lineages: subtypes of the intimin encoding gene *eae* and the cytolethal distending toxin (Cdt) gene *cdtB* were lineage associated, the second type three secretion system (ETT2) island was truncated in L3 and L6. Seven new *eae* subtypes and 1 new *cdtB* subtype (*cdtB*-VI) were found. Alarmingly, 85.9% of the Chinese *E. albertii* isolates were predicted to be multidrug resistant (MDR) with 35.9% harboured genes capable of conferring resistance to 10 to 14 different drug classes. By *in silico* multi-locus sequence typing, majority of the MDR isolates belonged to 4 STs (ST4638, ST4479, ST4633 and ST4488). Thirty-four intact plasmids carrying MDR and virulence genes, and 130 intact prophages were identified from 17 complete *E. albertii* genomes. Ten plasmid replicon types were found to be significantly associated with MDR. The 130 intact prophages were clustered into 5 groups, with group 5 prophages harbouring more virulence genes. Our findings provided fundamental insights into the population structure, virulence variation and MDR of *E. albertii*.

**Impact statement:** *E. albertii* is newly recognized foodborne pathogen causing diarrhea. Elucidation of its genomic features is important for surveillance and control of *E. albertii* infections. In this work, 169 *E. albertii* genomes from difference sources and regions in China were collected and sequenced, which contributed to the currently limited genomic data pool of *E. albertii*. In combination with 312 publicly available genomes from 14 additional countries, the population structure of *E. albertii* was defined. The presence and subtypes of virulence genes in different lineages were significantly different, indicating potential pathogenicity variation. Additionally, the presence of multidrug resistance (MDR) genes was alarmingly high in the Chinese dominated lineages. MDR associated STs and plasmid subtypes were identified, which could be used as sentinels for MDR surveillance. Moreover, the subtypes of plasmids and prophages were distributed differently across lineages, and were found to contribute to the acquisition of virulence and MDR genes of *E. albertii*. Altogether, this work reveals the diversity of *E. albertii* and characterized its genomic features in unprecedented detail.

**Data Summary:** All newly sequenced data in this work were deposited in National Center for Biotechnology Information (NCBI) under the BioProject of PRJNA693666, including 6 complete genomes and raw reads of 164 *E. albertii* isolates.

## Introduction

*Escherichia albertii* is a recently defined species and a recognised foodborne human pathogen [1–3]. *E. albertii* mainly causes diarrhea [3, 4], while bacteraemic human infections were also reported [5]. *E. albertii* has historically been misidentified as various pathogens such as enterohemorrhagic *Escherichia coli* (EHEC), enteropathogenic *E. coli* (EPEC), *Shigela boydii* serotype 13, and *Hafnia alvei* [1, 6]. In 2003, it was confirmed to be a novel species of the genus *Escherichia* and named as *E. albertii* [2, 6]. Through retrospectively studies, *E. albertii* was found to be responsible for a human diarrhea outbreak in Japan in 2011 [7]. *E. albertii* can also cause infections in other animals. An outbreak of *E. albertii* infection in common redpoll finches in Alaska led to deaths of hundreds of birds in 2004 [8]. Furthermore, *E. albertii* has also been isolated from a variety of sources including food products [9].

The pathogenicity of *E. albertii* was mainly attributed to a type III secretion system (T3SS) encoded by the locus of enterocyte effacement (LEE) and the cytolethal distending toxin (Cdt) encoded by the *cdtABC* operon, both of which were commonly found in *E. albertii* [1, 9, 10]. There were also multiple non-LEE effector genes [11]. Based on the presence of the intimin *eae* gene, the LEE locus was found to be widely present in *E. albertii* [1, 9]. The non-LEE effector genes, which were mainly acquired through prophages in *E. coli* [11], were observed in three *E. albetii* complete genomes [10]. Another *E. coli* type III secretion system 2 (ETT2), which has major effects on the surface proteins associated with motility and serum survival (as a prerequisite for bloodstream infections) of *E. coli*, has also been found in *E. albertii* [12]. ETT2 were predicted to be common in *E. albertii* based on the representative *eivG* gene [1, 10]. Shiga toxin (Stx) gene *stx2f* and *stx2a* are sporadically observed in *E. albertii* [1]. However, the detailed distribution of these genes in *E. albertii* remained unclear, and the other virulence factors reported in *E. coli* have not been systematically investigated in *E. albertii*.

Antimicrobial resistance (AR), especially multi drug resistance (MDR) which is defined as resistance to 3 or more drug classes, is an increasing global challenge [13]. Phenotypic AR and MDR of *E. albertii* strains were observed in Brazil and China, respectively [14, 15]. Poultry source *E. albertii* isolates in China were phenotypically resistant to up to 11 drug classes, some of which were commonly used in clinical treatment such as cephalosporins, aminoglycosides, fluroquinolones, and beta-lactam antibiotics [14]. However, the overall presence of AR genes in *E. albertii* isolates from different geographic regions and sources remains unclear.

It is well known that transmissible elements, especially plasmids and phages, are associated with the acquisition of virulence and AR genes [16]. They are key transmissible elements for the acquisition of *stx* genes, T3SS effector genes, and other virulence genes in *E. coli* [16]. Multiple intact plasmids of *E. albertii* carrying virulence and MDR genes were reported [1, 14, 17]. However, plasmids in draft genomes of *E. albertii* and their association with the acquisition of AR and virulence genes remain to be characterized [1, 10]. Prophages have been found in *E. albertii* with 4-7 prophages per genome from 3 complete genomes analysed [1]. However, their carriage of virulence and AR genes has not been examined.

Two clades of *E. albertii* have previously been defined based on whole genome sequencing analysis [1, 18], with no isolates from China. In this work, *E. albertii* from different sources and regions of China were isolated and sequenced, including 163 draft and 6 complete genomes. Publicly available complete genomes and draft genomes of *E. albertii* were analysed together to elucidate the population structure, virulence and resistance of *E. albertii* and the relationships of Chinese and international isolates.

## Methods

### Genomic sequences

A total of 169 *E. albertii* isolates from different sources and regions in China were collected and sequenced. The *E. albertii* type strain LMG20976 was also sequenced in this study. All of the isolates were sequenced using Illumina sequencing [19], except for 6 isolates that were additionally sequenced using Pacbio [20] to obtain complete genomes.

Raw reads and assemblies of publicly available *E. albertii* isolates were downloaded. To identify *E. albertii* isolates that were potentially misidentified as *E. coli*, one reported specific gene (EAKF1_ch4033) of *E. albertii* [21], was searched against a total of 29,988 *E. coli* (including *Shigella)* genome assemblies using BLASTN, with the thresholds of coverage 50% and identity of 70%.

In summary, there were a total of 482 genomic sequences of *E. albertii* included in this study (**Table S1**). For draft genome sequences, 164 were from this study and 296 were from public databases (255 raw reads from European Nucleotide Archive and 41 assemblies from NCBI). For complete genomes, there were 6 from this study, and 16 genomes from NCBI (10 of which were sequenced by PacBio). Raw reads of Illumina sequencing were assembled using Skesa v2.4.0 [22].

### Phylogenetic analysis and in silico multi-locus sequence typing (MLST) of *E. albertii*

In an initial analysis, 38 representative isolates were selected to represent *E. albertii* diversity to obtain the over picture and to identify the root of the *E. albertii* phylogeny. Using *E. coli* (Accession No. NZ_CP014583.1) as reference, SNPs were called by snippy v4.4.0 [23], and recombinant SNPs were detected and removed by Gubbins v2.0.0 [24]. A maximum parsimony tree based on SNPs of the 38 isolates using *E. coli* as outgroup was constructed by Mega X with 1000 bootstraps [25].

To elucidate the phylogenetic relationship of the 482 *E. albertii* isolates, a phylogenetic tree was constructed using SaRTree v1.2.2 with ASM287245v1 as reference [26]. The recombination sites of the SNPs were removed using Recdetect v6.0 [26]. The SNP alignment of the genomes were analysed with Fastbaps v1.0.4 to identify lineages of *E. albertii* [27]. The lineages defined were mapped onto the phylogenetic tree using ITOL v4 [28].

The *in silico* MLST based on the 7 housekeeping genes of *E. coli*, were performed on *E. albertii* with sequence types (STs) assigned [23, 29]. Clonal complexes (CCs) of the STs were called based on one allele difference using the eBURST algorithm [39].

### Virulence and antibiotic resistance analysis of *E. albertii*

Predicted virulence and antimicrobial resistant genes from the *E. albertii* genomes were identified by Abricate v0.8.13 [23]: Virulence genes were screened against the *E. coli* virulence factors database (Ecoli_VF) and the virulence factor database (VFDB) with identity of >= 70% and coverage of >= 50% [30]; Antibiotic resistant genes were screened through the NCBI AMRFinder database with identity of >= 90% and coverage of >= 90% [13];

To predict the subtypes of the *eae* and *cdtB* genes harboured by each *E. albertii* isolate, representative sequences for each type of *eae* and *cdtB* were used to search the collection of *E. albertii* genomes using BLASTN with identity of >= 97% and coverage of >= 50% [31]. The new *eae* and *cdtB* subtypes were defined based on the tree structure and BLASTN results. A new subtype was defined, if it was phylogenetically distant from the known subtypes and was present in >= 5 isolates (with identity >= 97%). The detailed methods for single gene phylogenetic tree construction for *eae* and *cdtB* were described in **supplementary methods**.

### Plasmid and prophage analysis based on complete genomes of *E. albertii*

For intact plasmids and prophages of *E. albertii*, 16 complete genomes by PacBio and one reference genome GCA_001549955.1 (sequenced by 454 GS-FLX) were selected for the prophage and plasmid analysis.

To identify the plasmids in the draft genomes, we used both PlasmidFinder and MOB-suite [23, 32]. Plasmid replicon genes were screened against the PlasmidFinder database with identity of >= 50% and coverage of >= 50% using Abricate v0.8.13 [23]. MOB suite was able to identify the potential plasmid sequences in draft genomes. MOB types were assigned if the predicted plasmids were known. To evaluate if the presence of the invasive plasmid pINV of *Shigella* present in *E. albertii*, the pINV specific gene *ipaH* and 39 plasmid-borne virulence genes were screened in the raw reads of *E. albertii* using ShigEiFinder [33]. AR genes and virulence genes present on the intact plasmids and MOB suite predicted plasmids were screened using the aforementioned criteria.

The complete genomes were submitted to Phaster for prophage prediction [34]. In order to define the groups of the intact prophages, the genomic sequences of prophages were annotated with Prokka v1.12 [35]. The gff files of the intact prophages were clustered by Roary v3.11.2 with identity of >= 70%, and a binary gene presence and absence tree was generated [36]. The concatenated prophage sequences in the order of binary clustering were visualized in similarity plots by Gepard v1.40 [37]. Genes whose presence was significantly associated with prophage groups (P <= 0.001) were identified using Scoary [38]. The top 3 to 5 genes that are of 100% specificity and sensitivity for each prophage group were identified as potential prophage specific genes. These prophage specific gene candidates were searched against the 482 genomes with identity >= 70% and coverage >= 50% using BLASTN. The distribution of the prophage specific genes were visualized in Phandango [39]. AR genes, plasmid replicon genes and virulence genes present on the intact prophages were screened using the aforementioned criteria.

To compare the prophages of *E. albertii* with public phage clusters from the Microbe Versus Phage (MVP) database, the representative phage sequences of different phage clusters were downloaded [40]. Each prophage sequence of *E. albertii* was searched against the MVP reference phage cluster sequences with identity of 80% and coverage of 50% using BLASTN [40].

## Results

### A dataset representing *E. albertii distribution* in different source types and geographic regions

A total of 169 *eae* gene positive *E. albertii* isolates from different regions of China were collected from 2014 to 2019 and sequenced in this study. The *E. albertii* isolates were from five provinces in China, the majority of which were from Sichuan province in Southern China and Shandong province in Northern China (**Table S1**). The Chinese *E. albertii* isolates belonged to 7 different source types, with 90.5% from poultry intestine (with 110 isolates from chicken intestines and 43 from duck intestines). There were 6 human source isolates from China (**Table S2**). Three isolates were from patients with diarrhea, including one patient with bloody diarrhea. Three *E. albertii* isolates were from poultry butchers and retailers who were asymptomatic. Two *E. albertii* isolates were from the faecal samples of bats in Yunnan, China. Notably, as only *eae* positive samples were cultured for *E. albertii* in this study, any *eae* negative *E. albertii* isolates would have not been isolated.

To compare the genomic characteristics of *E. albertii* globally, a total of 312 publicly available *E. albertii* genome sequences were included in this study. Based on the metadata available, these isolates were from 6 continents and 12 different source types including humans, birds, bovine, swine, cats, water mammals, camelid, plants, soil and water. Humans (76 isolates) and birds (30 isolates) were the dominant sources (**Table S3**).

All 482 genomes were screened using the *E. albertii* specific gene marker (EAKF1_ch4033) [21] with 4 isolates being negative. Phylogenetic analysis confirmed the 4 EAKF1_ch4033-negative isolates belonged to the *E. albertii* clade 1 as described below.

### *E. albertii* lineages and their distribution in different geographic regions and source types

Previous studies showed that *E. albertii* is divided into 2 clades [1, 18]. To better define the phylogenetic lineages, we used Fastbaps to analyse the population divisions of the 482 *E. albertii* isolates using non-recombinant SNPs (with recombinant SNPs removed) as input. Eight lineages of *E. albertii* were defined (353 isolates) while 129 did not belong to any lineage (**Figure 1**) [27]. Lineage 1 (L1) corresponds to previously defined clade 1, and L2 to L7 belonged to the previously defined clade 2 [1, 18]. It is noteworthy that the *E. albertii* isolates which were previously identified as *S. boydii* serotype 13 belonged to L3. Each lineage includes isolates from multiple continents. L5 and L8 were more common in Asia, while L1 (or clade 1), L3 and L6 were more common in Europe and North America (**Figure S1**).

**Figure 1.**
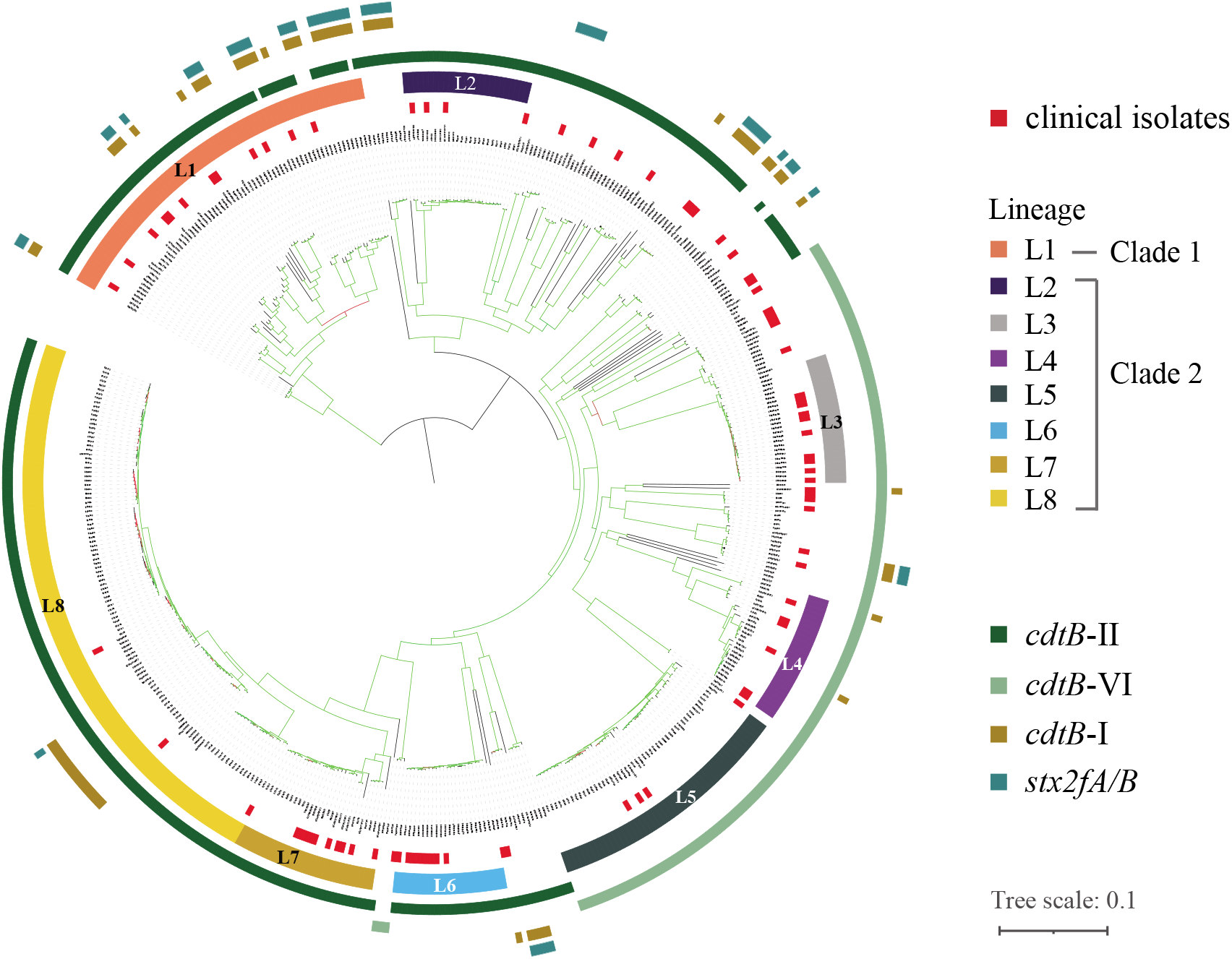
Phylogenetic structure of *E. albertii*. The phylogenetic tree of the 482 *E. albertii* isolates was constructed using Quicktree with bootstraps of 1000 [26]. The colour of the branches represented the percentage of bootstrap supporting from 10% to 100% (from red to green). The inner most ring marks the isolates from human clinical source. The next ring marks the lineages by colour as shown in the colour legend. The outer 4 rings represented the *cdtB* subtypes and the *stx*2f gene, which were represented with different colours as shown in the colour legend.

The 85 human clinical isolates were distributed among the 8 lineages indicating all of these lineages were potentially pathogenic to humans (**Figure 1**). For Chinese *E. albertii* isolates, the 6 human clinical isolates belonged to L4 (2), L7 (1), L8 (1), with two not falling into any lineages (**Table S2**). The two bat source isolates did not belong to any of the lineages but were most related to L3. There were 158 poultry source isolates from China, 55.7% of which belonged to L8 followed by L5 (22.8%) (**Table S3**), and there were two isolates of L8 from wild birds. By contrast, the majority of the bird source isolates from other countries came from wild birds, 53.3% of which did not belong to any of the 8 lineages while 33.3% were from L1. These findings demonstrated that the bird source *E. albertii* isolates from the other countries were phylogenetically different from the wild birds and poultry source isolates in China.

### In silico MLST of *E. albertii* isolates

We performed *in silico* MLST on the isolates using the established *E. coli* scheme [29]. The 482 *E. albertii* isolates were subtyped into 98 STs, among which 53 STs contained >= 2 isolates. By lineage, with the exception of L1 and L8, each lineage was dominated by one ST. ST4633 accounted for 84.0% of the total number of isolates in L2, ST5431 for 76.0% of L3, ST4619 for 60.0% of L4, ST4638 for 81.3% of L5, ST5390 for 100% of L6 and ST3762 for 82.1% of L7. And 94.6% of L8 belonged to 4 STs (ST4488, ST4634, ST4479 and ST4606). We further grouped closely related STs as CC using one allele difference [41]. Nearly half of the STs (43 of 98) were grouped into 9 CCs while the remaining 55 STs were singletons (**Figure 2A**). With the exception of L4 and L6 which only contained STs, the other lineages were dominated by one CC. CC1 represented 68.1% of the L1 isolates. CC2 to CC6 were representative of more than 90% of the isolates in L2, L3, L5, L7 and L8 respectively. The majority of the singletons (42 of 55) belonged to none of the 8 lineages and were classified as other in the lineage division above.

**Figure 2.**
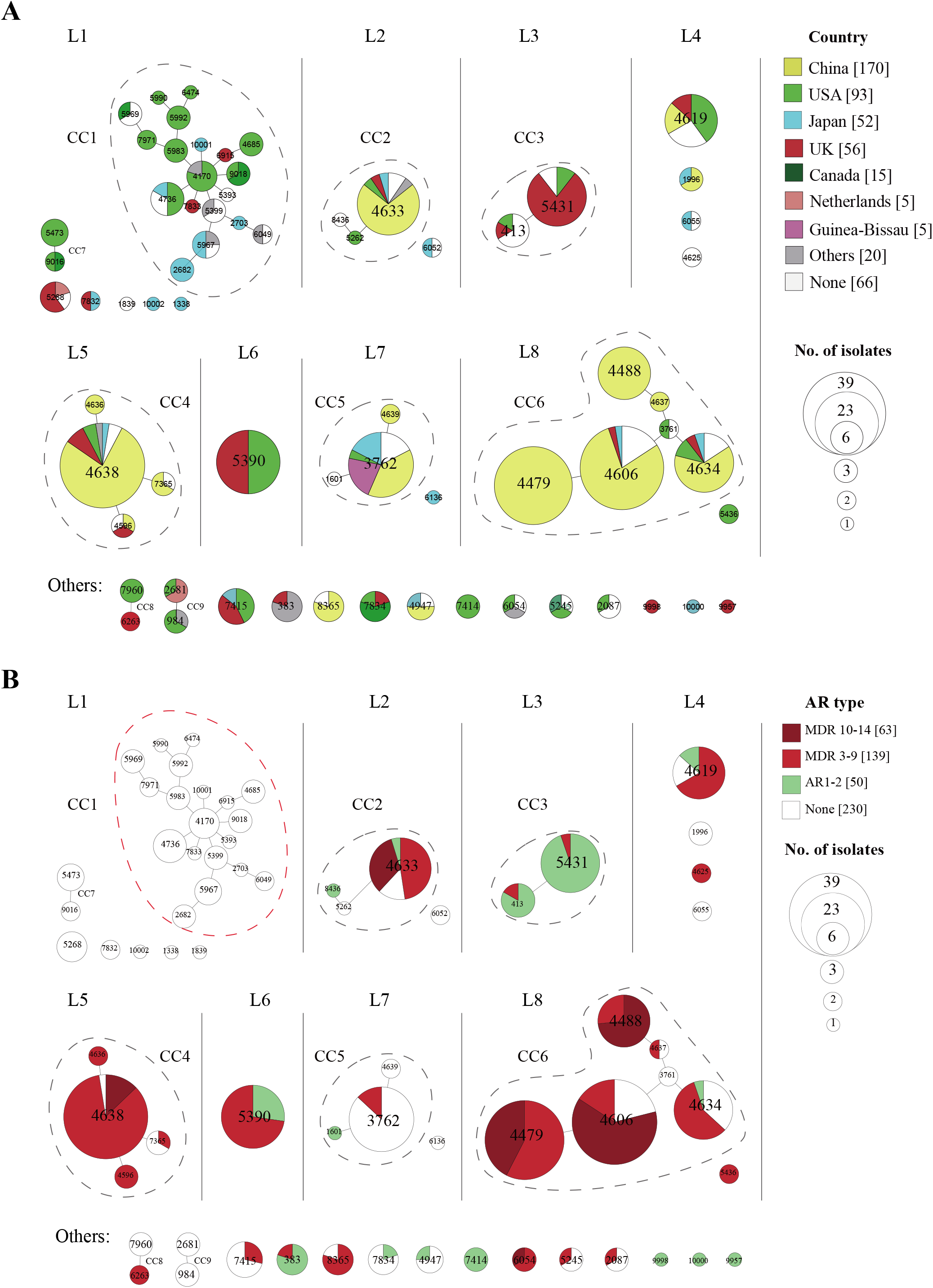
Region distribution and resistance profiles of clonal complex (CC) and sequence type (ST) of *E. albertii* isolates based on the 7 gene multi-locus sequence typing (MLST). (A). Region distribution of STs and CCs. (B). Drug resistance profiles of STs and CCs. Each circle represented an ST and the size of the circles reflected the number of isolations. STs and CCs belonging to different lineages were separated. STs with one allele difference were linked with solid lines as one CC. Singleton STs were shown for each lineage. While for the 42 singleton STs belonging to none of the 8 lineages, only 12 STs with AR genes were shown. The top 7 countries with 5 or more isolates were highlighted in different colours as shown in the colour legend. Antibiotic resistance of different STs is denoted by different colours of different level of resistance as shown in the colour legend. The pie chart within an ST denotes of different proportions of isolates displaying a particular characteristic.

Thirty-three STs were found in more than one country while 57 STs were only found in one country. The six largest CCs were found in more than one country. However, individual STs or CCs were predominant in different countries or regions. ST5390 was the most common ST in both USA and UK, and ST5431 was the second most common ST in the UK. In China, ST4479, ST4638 and ST4606 were the main STs, representing 54.7% of the Chinese isolates. CC1 and CC3 were predominant in the USA and UK while CC2, CC4, and CC6 were predominantly found in China.

### Virulence genes and their distribution in *E. albertii* lineages

Virulence genes from *E. coli*_VF database were screened to evaluate the potential pathogenicity of *E. albertii*. The LEE island from LEE1 to LEE7 contains 41 genes [42]. The 41 genes were present in slightly different proportions ranging from 91.1% to 99.8%, with the *espF* gene the lowest in 439 of the 482 isolates (**Table S4**). The *eae* gene on LEE5 was harboured by 99.4% (479/482) of the isolates. Thirteen previously defined *eae* subtypes were observed in 387 (80.3%) of the 482 isolates, and 7 new *eae* subtypes were identified (which were observed in >= 5 isolates each) among the remaining 92 isolates (**Figure S2A**). Subtype sigma was the dominant type (37.9%), followed by rho (10.4%), itota2 (6.6%) and epsilon3 (6.2%) (**Figure S2B**). The *eae* subtypes were associated with specific lineages: epsilon3, iota2 and rho were the predominant subtypes in L2, L3, L5 respectively, and subtype sigma was dominant in L6, L7 and L8. However, L1, L4, L5 and L7 harboured multiple *eae* subtypes. L1 (or clade 1), possessed 8 *eae* subtypes, with beta3, alpha8 and the newly defined sigma2 and alpha9 as the main subtypes (**Figure S2C**).

Cdt facilitates bacterial survival and enhances pathogenicity [43] and is encoded by the *cdtABC* genes which were widely distributed in *E. albertii* [1, 44]. In this study, *cdtABC* genes were present in 99.4% (479/482) of the isolates. The *cdtB* gene had been previously divided into five subtypes (*cdtB*-I to *cdtB*-V), with *cdtB*-II/III/V as one group, and *cdtB*-I/IV as another group [45]. By phylogenetic analysis of the *cdtB* genes in *E. albertii*, a new *cdtB* subtype was identified and named as *cdtB*-VI. *E. albertii cdtB*-VI was phylogenetically closer to *cdtB* group II/III/V (**Figure S3**). Notably, almost all *cdtB*-VI positive *E. albertii* isolates (30.1%, 145/482) were located on the same branch that includes L3, L4 and L5 isolates (**Figure 1**). *CdtB*-II, as the dominant type, was present in 68.3% (329/482) of *E. albertii* isolates across 5 lineages (L1, L2, L6, L7 and L8)*. CdtB*-I was found in 65 (13.5%) *E. albertii* isolates, 89.2% (58/65) of which were also positive for either *cdtB*-II or VI. There were 49 isolates positive for *sxt2f* (10.2%, 49/482), 44 of which possessed *cdtB*-I (**Figure 1**). *E. albertii* isolates with *cdtB*-I were significantly more likely to harbour *sxt2f* gene (Chi-Square test, P<0.001). Both *cdtB-I* and *stx2f* were observed on the same intact prophage of two complete genomes (ASM331252v2_PF4 and ASM386038v1_PF5). None of the Chinese *E. albertii* isolates were positive for *stx2f*.

ETT2, which plays a role in motility and serum resistance in *E. coli* [12], was found to be nearly intact in 61.4% (296/482) of the isolates, except for the *ygeF* gene which was absent in all *E. albertii* isolates [10]. Eighty-eight isolates (18.3%) harboured 29 to 31 ETT2 genes with 2 to 4 genes missing. Interestingly, ETT2 genes were mostly deleted in L3 and L6 with only 4 and 3 genes remaining, respectively (**Figure 3**). Other virulence genes were also lineage restricted such as the type VI secretion system (T6SS) *aec* genes, which were present in most of the lineages except L1, L3 and L5. The haemolysin genes *hlyABCD* were present only in L3 isolates (**Figure 3**). The *iuc* gene cluster (*iuc-ABCD* and *iutA*) which encodes aerobactin [46] was mainly present in L3, L4 and one isolate of L6. The *Yersinia* high pathogenicity island (HPI), which encodes the yersiniabactin (Ybt) [47], was only found in L6 isolates (100%). The *lng* gene cluster that encodes the CS21 pilus (class b type IV) [48–50] was mainly observed in L5.

**Figure 3.**
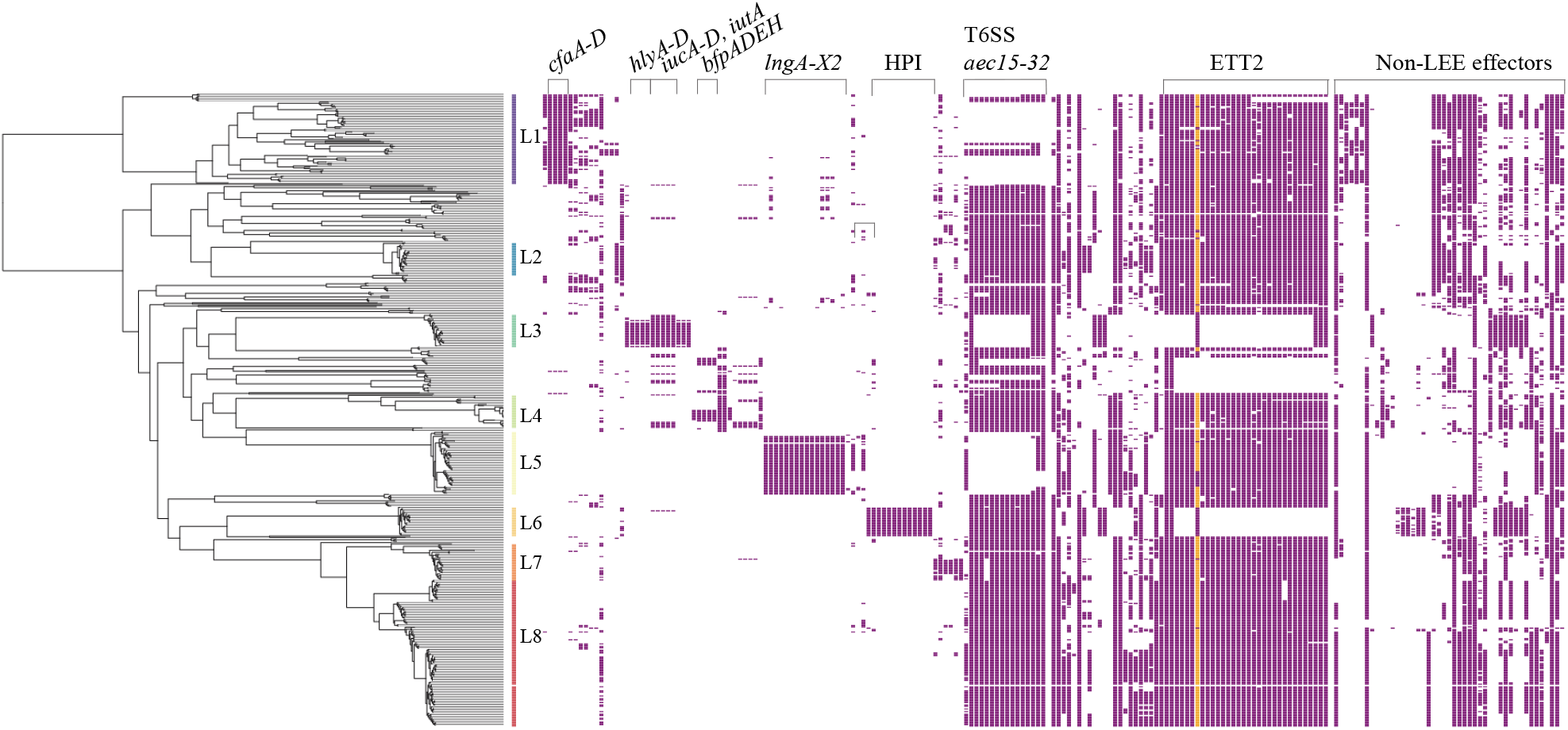
Virulence genes that were significantly associated with different lineages of *E. albertii*. The distribution of different virulence genes in *E. albertii* were visualized using Phandango [39]. The lineages of *E. albertii* were labelled with different colours. The presence of a gene was marked with a coloured box. Only genes or gene clusters significantly associated with lineages are shown.

There were other *E. coli* virulence genes including *paa, efa1*, the bundle forming pilus (BFP) encoding *bfp* genes that were found to be variably present in *E. albertii* which are summarised in **Table S4**. One genome assembly (ERR1953722) from L5 was found to harbour *Shigella* invasive plasmid pINV genes [51]. However, further investigation by read mapping found that it was most likely due to contamination (data not shown).

### Drug resistance genes and their high prevalence in some STs of *E. albertii*

Presence of AR genes was screened using NCBI AMRFinder database [13]. Among the 482 isolates, 52.3% (252/482) harboured AR genes, 41.9% (202/482) were MDR (harbouring AR genes resistant to >= 3 different drug classes), and 13.1% (63/482) of the isolates harboured genes capable of conferring resistance to 10 to 14 different drug classes that were regarded as highly resistant. Notably, 72.3% (146/202) of the predicted MDR isolates were from China with AR rate of 88.2% and MDR rate of 85.9% with 61 isolates (35.9%) being predicted to be highly resistant. The predicted AR drug classes were shown in **Figure 4**, including sulfamethoxazole-trimethoprim, cephalosporin, streptomycin, beta-lactam antibiotics, etc. The antibiotic resistance genes observed in each isolate were shown in **Table S5**.

**Figure 4.**
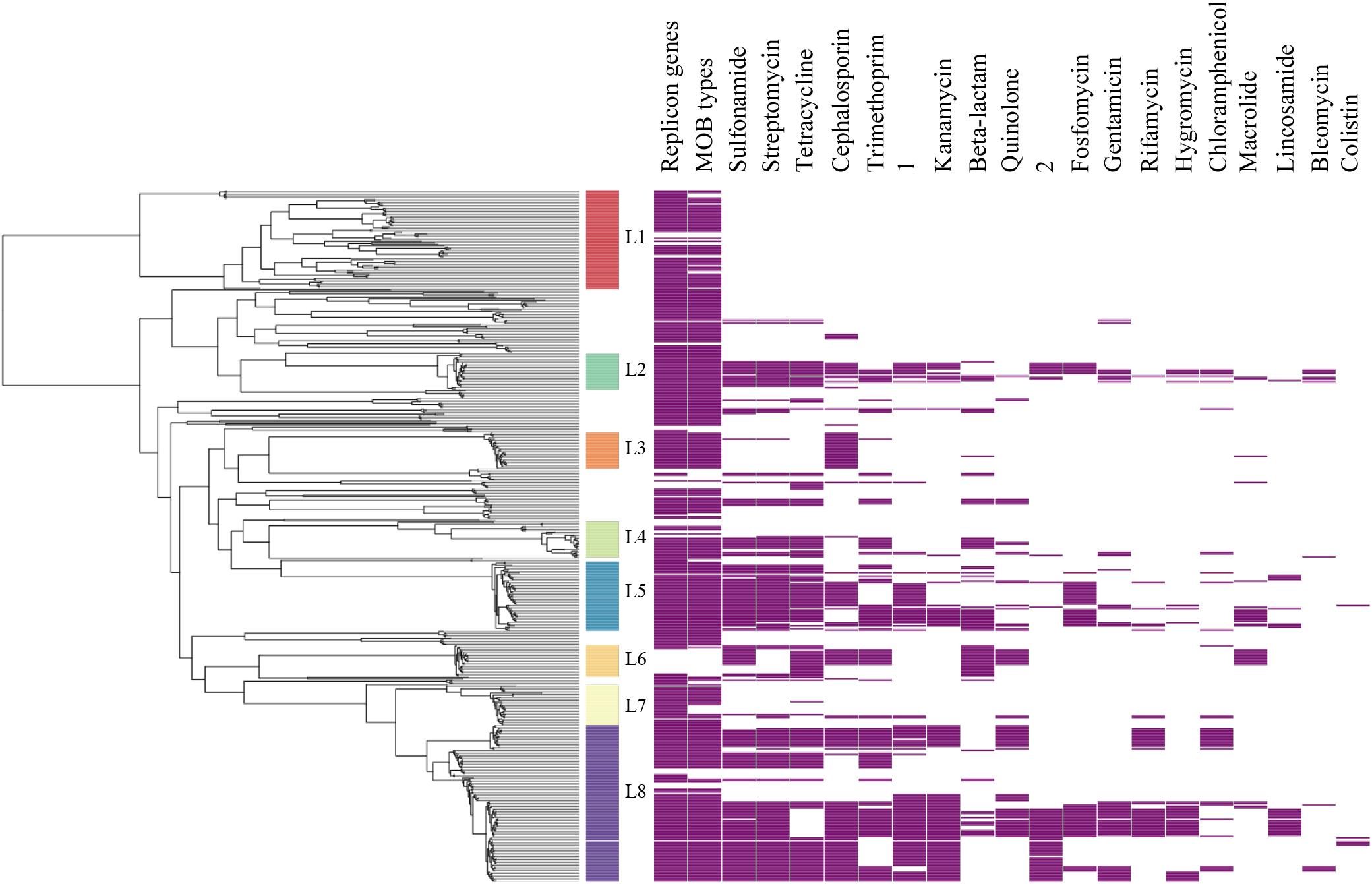
Predicted resistance to drug classes in *E. albertii*. *E. albertii* isolates that harboured genes conferring resistance to different drug classes are shown in purple. The two columns headed with 1 and 2 denote combination of 2 drugs as follows: 1 = chloramphenicol and florfenicol, 2 = phenicol and quinolone. Isolates with predicted plasmids by PlasmidFinder and MOB suite (respectively) were also highlighted.

We determined resistance profiles by STs and found that some STs contained a high proportion of MDR isolates. The predicted MDR rates in ST4638, ST4479, ST4633 and ST4488 were >= 80% (**Figure 2B**). Additionally, 63.2% of the isolates in ST4606 were highly resistant. For the top 6 STs in China representing 84.7% (144/170) of the Chinese isolates, 94.8% (135/144) of the isolates were predicted to be MDR, and 41.7% (60/144) were highly resistant. In contrast, isolates from the USA and UK had relatively lower predicted MDR rate (26.2%, 39/149) and were mainly observed in ST5390, ST4619 and ST4638, with only one highly resistant isolate (**Figure 2**). By CCs, CC3, CC4 and CC6 had high MDR rate. CC1 carried hardly any resistance genes while CC3 and CC5 had low levels of carriage of resistance genes.

### Plasmids and plasmid associated drug resistance and virulence genes

We firstly analysed the 17 complete *E. albertii* genomes for the carriage of plasmids. There were 34 intact plasmids ranging from 19,118 bp to 265,919 bp (**Table S6**). Nineteen plasmids were previously reported [1, 14, 17], while 15 plasmids were newly identified in this study.

We further performed plasmid typing using PlasmidFinder and MOB-suite [23, 32]. PlasmidFinder identifies plasmid by replicon types [23]. However, it should be noted that a plasmid may carry more than one replicon type. MOB-suite predicts plasmid using the relaxase gene and group those predicted plasmids into different MOB types [32]. However, some plasmids have no relaxase genes. Thus, both methods were used to predict and identify plasmids in all *E. albetii* isolates. Among the 482 *E. albertii* isolates, PlasmidFinder found that 86.7% (418/482) of the isolates harboured plasmids, with a total of 54 replicon types detected. There were 34 replicon types that each was present in more than 10 isolates. And 26 replicon types were found to be significantly associated with lineages (P < 0.001) (**Table S7**): for example, IncFII(29)_1_pUTI89 type with L2, Col156_1 with L3, and IncFII (pSE11)_1 with L4, IncX1_1 with L5 and L8. By MOB-suite, a total of 1854 plasmid sequences were predicted in 427 of the 482 isolates with an average of 4.3 plasmids per genome while 55 isolates had no plasmids predicted. The vast majority (90.3%, 1674/1854) of the predicted plasmids were grouped into 170 MOB types with the remaining 9.7% (180/1854) being novel with no MOB types. There were 47 MOB types each of which was present in >= 10 isolates, 36 of which were significantly associated with lineages, which is concordant with findings from replicon types (**Table S7**). Additionally, there were 64 isolates without both replicon types and MOB types observed, including 77.3% (17/22) of L6 isolates (**Figure 3**). However, 35.9% (23/64) of these isolates harboured AR genes, especially 72.3% of L6 were predicted to be MDR.

Plasmids are known to be responsible for the acquisition of MDR genes. Among the 34 intact plasmids, 9 were found to harbour AR genes (**Table S6**). One newly identified MDR plasmid, ESA136_plas1 (MOB type AA738), which contained 15 AR genes resistant against 13 drug classes, harboured IncHI2_1, IncHI2A_1 and RepA_1 replicon types.

Statistical association between MDR and the plasmid types were evaluated. By PlasmidFinder, 13 replicon types were found significantly associated with MDR (P < 0.001, Chi-square test) (**Figure S4**). However, this analysis may be biased when the MDR genes were not located on the same plasmid with the replicon genes. This bias can be resolved by MOB-suite, which offers the predicted plasmid sequences from the draft genomes. We screened the plasmid replicon genes and MDR gene on the MOB-suite predicted plasmids. Ten replicon types were confirmed to be significantly more likely to be observed in MDR isolates (P < 0.001) including IncQ_1, IncN_1, ColE10_1, IncHI2A_1, RepA_1, IncHI2_1, IncFII(pSE11)_1, IncX9_1, IncFII(pHN7A8)_1, and IncX1_1. The predicted odds ratio (OR) values ranged from 6.1 to infinity (**Figure 5A**). Further, each MOB type possessed 1 to 8 plasmid replicon genes, indicating MOB typing is of higher resolution than replicon typing (**Table S7**). Five MOB types AE928, AA860, AA738, AA334 and AA327 were significantly associated with MDR genes (P<0.001, OR 15.0 to infinity) (**Figure 5B**). Importantly, the MDR associated replicon types and MOB types were mainly observed in L4, L5 and L8, which had a high proportion of MDR isolates.

**Figure 5.**
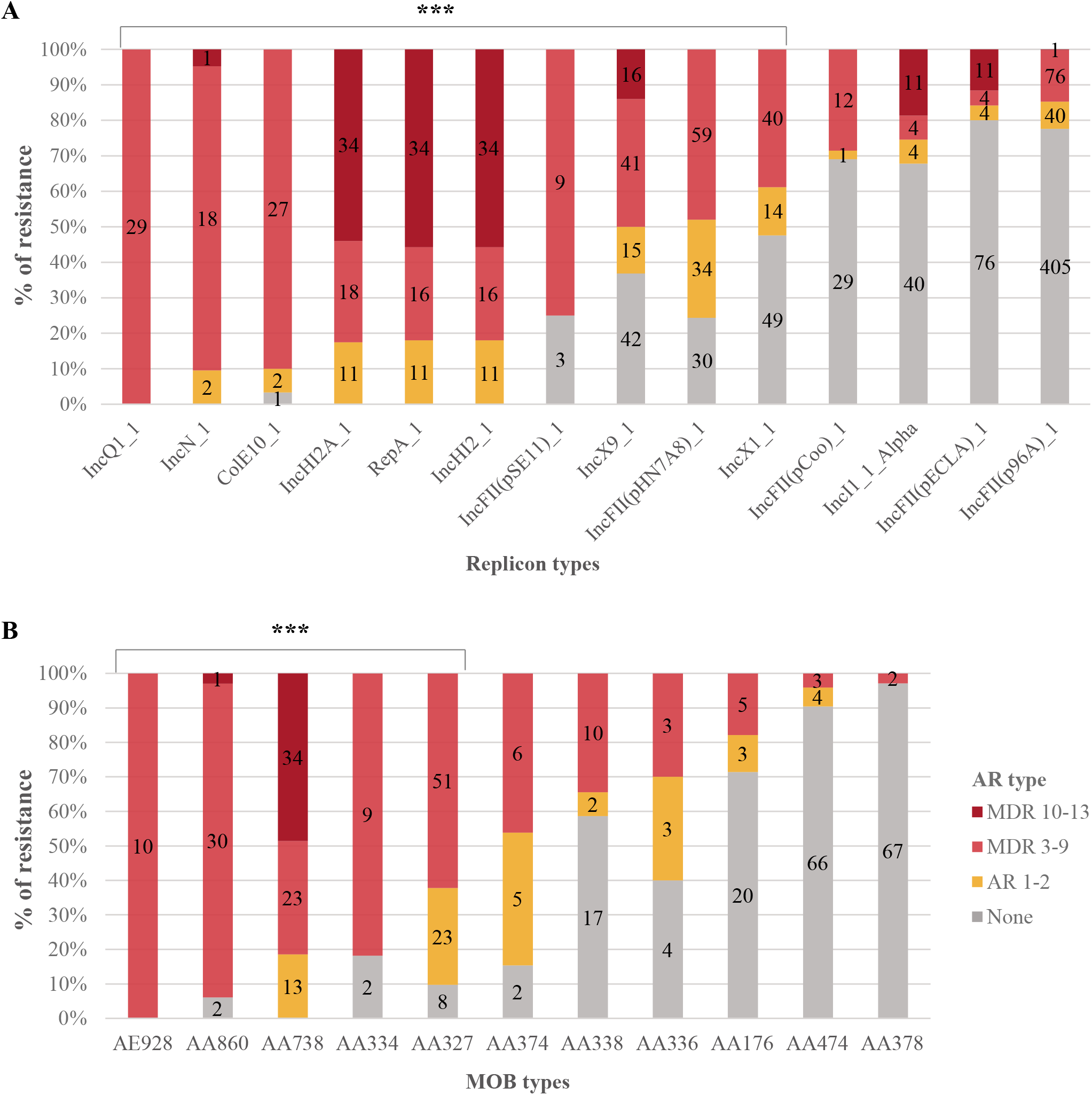
Multidrug resistance (MDR) associated plasmid subtypes. (A). Replicon types detected. (B). MOB types detected. Those types significantly associated with MDR are marked with P value < 0.001 (***). The proportion of drug resistance (%) for each replicon or MOB type was shown as colour legend.

Lastly, association of virulence genes with plasmids were evaluated. Among the 34 intact plasmids, 27 harboured virulence genes. Two plasmids from bat source isolates harboured the Type II secretion system and the putative heat-stable enterotoxin gene *astA* [52] (**Table S6**). Moreover, some lineage restricted virulence genes were observed in the MOB suite predicted plasmids, including the *LngA-lngX* gene cluster, the *iucA-iucD* gene cluster, and the *hlyABCD* gene cluster.

### Prophages and carriage of resistance and virulence genes

PHASTER was used to search for prophages in the 17 complete genomes first [34]. A total of 207 prophages were identified: 130 were intact, 50 were incomplete and 27 were indeterminant (**Table S9**). The size of the intact prophage genomes ranged from 11.163 to 98.311 kb. Most of the intact prophages were integrated on the chromosomes with 11 (8.5%) being on plasmids.

We grouped the 130 intact prophages based on a tree generated using the presence/absence of prophage genes using Roary v3.11.2 [36], and a nucleotide dotplot generated using Gepard v1.3 [37]. Gepard was a useful method for grouping diverse prophages [53]. As seen in **Figure 6**, the darker the colour in the dotplot, the more similar the sequences were. There were 5 main squares with dense dots corresponding to 5 main groups of prophages (G1-G5). G5 was more diverse and potentially can be further subdivided subgroups. Of prophages in G1 and G2, 50% (4/8) and 85.7% (6/7) (respectively) were from the two bat source isolates.

**Figure 6.**
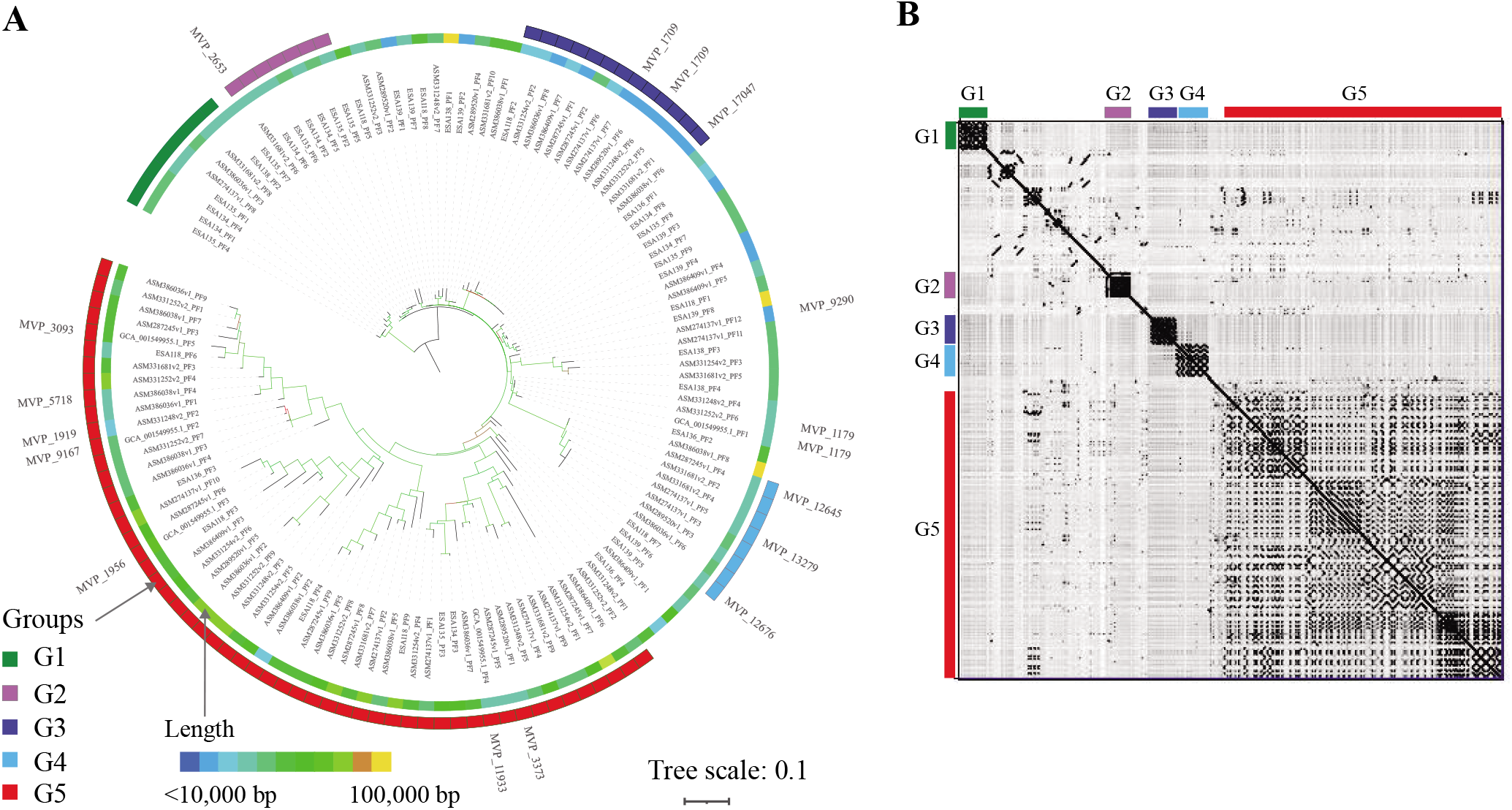
Clustering of the intact prophages of *E. abertii*. (A). Accessory binary gene presence tree of the prophages constructed using Roary v3.11.2 [36]. The 5 main groups of prophages were labelled with different strip colours. There were 15 prophages of *E. albertii* with phage cluster types in the Microbe Versus Phage (MVP) database, the 15 MVP phage cluster types were labelled. (B). Dot plot of similarity of prophages using the nucleotide dotplot tool GEPARD [37] and the 5 prophage groups were marked.

Based on the annotation of the 130 intact prophages, genes that were present only in one prophage group were identified using Scoary [38], and were designated as group specific gene markers for each of the prophage groups. By screening the group specific genes among the draft genomes, G1 was predicted be present in 34.4% (166/482) of the *E. albertii* isolates, with at least two specific genes of G1 identified in these genomes. G2 was predicted to be in 3.7%, G3 in 46.7%, G4 in 59.1% and G5 in 96.1% of the 482 *E. albertii* isolates (**Figure S5**). In terms of lineage distribution, G3 prophage specific genes were more likely to be observed in L5 and L8, and G4 prophages in L3, L4 and L8 (P < 0.001, OR value > 3.9). G1 prophage specific genes were negatively associated with L3 and L6, G3 prophages with L2, L3, L6 and L7, and G4 prophages with L2, L5, L6 and L7 (P < 0.001, OR value < 0).

There were 27 T3SS non-LEE effector genes present in 59 of the 130 intact prophages, 64.7% of which were in G5 prophages (**Table S9**). Two intact G5 prophages were positive for both *stx2f* and *cdtABC* genes. Additionally, there were 3 intact prophages harbouring AR genes and all 3 were located on plasmids.

The MVP database collected viral genomes and prophage sequences from bacterial and archaeal genomes [40]. Those virus and prophage genomes were clustered based on their sequence similarity, with unified cluster types assigned [40]. By nucleotide comparison with the MVP representative phage clusters database using BLASTN, only 13.1% (17/130) of the intact prophage sequences were previously recorded in the MVP database, belonging to 15 phage cluster types (**Figure 6A**), indicating high diversity of prophages in *E. albertii* which have not been recorded in the database. Interspecies transmissions of prophages were observed: among the 15 MVP phage clusters, 11 prophages were previously observed in *E. coli*; cluster 12645 was previously observed in both *E. coli* and *Salmonella enterica*; and cluster 17047 from *Salmonella enterica*, while 5 phage clusters were only observed in *E. albertii*. In the 5 groups of prophages, MVP phage clusters were observed in G1, G3, G4 and G5, indicating G2 is a new prophage group specific for *E. albertii*.

## Discussion

*E. albertii* is a newly defined species of *Escherichia*, with infections previously wrongly attributed to *E. coli* and *Shigella* owing to the lack of sufficient subtyping techniques [1, 2, 18]. The *eae* gene and *cdtB* gene have since been used for *E. albertii* identification [9, 21, 54]. However, both genes were not present in all *E. albertii* isolates or unique to *E. albertii*. In this work, only *eae* positive samples were cultured for *E. albertii*, which would have missed any potential *eae* negative *E. albertii* isolates.

Previous study defined two clades of *E. albertii*, which was supported by this study [18]. Further, a total of 8 robust lineages were defined in this study. Clade 1 corresponds to L1, and clade 2 was further divided into 7 lineages (L2 to L8). The genomic features of these lineages were characterized. Based on the 7 gene MLST of *E. coli* [29], lineage representative STs (e.g. ST4638 for L5 and ST5390 L6) and CCs were identified. The stable and unified nomenclature characteristics of STs are more efficient in the global surveillance system [55]. Thus, using STs or CCs as hallmarks for different lineages of *E. albertii* will be useful when genomic information is not available, which would facilitate comparison between different studies and surveillance of global spread and MDR. Although the isolates sequenced may not be representative, lineages were of significantly different proportions in different geographic regions: L5 (represented by ST4638) and L8 (represented by 4 STs) were more common in China, and L3 and L6 were only observed in Europe and North America. This study showed the high diversity of *E. albertii*, and more lineages are likely to be identified with more isolates sequenced. Isolates causing human infection were observed in all 8 lineages, indicating all lineages are potentially pathogenic.

### Virulence gene variation in different lineages of *E. albertii*

The T3SS and the Cdt are the main virulence factors present in the vast majority of the *E. albertii* isolates. However, the subtypes of *eae* and *cdtB* were phylogenetically diverse. The *eae* gene was more diverse than the *cdtB* gene, and different lineages were dominated by different *eae* subtypes. Thus, it is likely that multiple independent acquisitions of the *eae* subtypes have occurred in *E. albertii*. There were 7 new *eae* subtypes identified, and these *eae* subtypes were phylogenetically distant from each other, indicating potential independent acquisition. It is also possible that these new *eae* subtypes evolved within *E. albertii*. For the *cdtB* gene, *cdtB*-II was dominant and present in all lineages except L3, L4 and L5 whereas the newly defined *cdtB*-VI was found in L3, L4 and L5. Given the phylogenetic relationship of the lineages, *cdtB*-VI must have replaced *cdtB*-II in L3-L5. However, it is unclear if the *cdtB*-VI evolved within *E. albertii* or was acquired from other species. Moreover, some subtypes of *eae* and *cdtB* were prevalent in *E. coli* but were rare in *E. albertii* and vice versa. For example, *cdtB*-III and V were common in Shiga toxin-producing *E. coli* (STEC), but were not observed in *E. albertii* [44, 56]; the *E. coli* prevalent *eae* subtypes were not common in *E. albertii* [57]; and the *eae* iota2 was observed in *S. boydii* serovar 13 isolates, which are in fact *E. albertii* [58]. The *eae* and *cdt* virulence genes seemed to have been acquired by *E. albertii* multiple times during its long evolutionary history. More studies are required to elucidate the interspecies and inter-species transfer of *eae* and *cdt* genes in the genus *Escherichia*.

Some virulence genes and pathogenicity islands were found to be associated with certain lineages. ETT2, which contributes to motility and serum resistance (which is essential for the invasive infections) in *E. coli* [12], was truncated in L3 and L6, while in the other lineages only the *yqeF* gene of ETT2 was absent. Experimental evaluation is required to determine whether ETT2 is functional without the *yqeF* gene in *E. albertii. Yersinia* HPI encodes the siderophore yersiniabactin (Ybt) for iron scavenging, which causes oxidative stress in host cells and contributes to the invasive extra-intestinal infections [47]. HPI comprises 11 genes, all of which were only observed in L6 isolates of *E. albertii*. Moreover, the *iuc* gene cluster include the *iucABCD* encoded the siderophore aerobactin and the *iutA* encoded ferric aerobactin were also associated with iron acquisition [46, 47]. The *iuc* gene cluster was mainly present in L3, L4 and one isolate of L6. More studies are required to evaluate the pathogenicity of those lineages that were equipped with different iron uptake systems. There were other lineage restricted virulence genes like T6SS, *hlyABCD* and the *lng* gene cluster. Although their expression remains unknown, these lineage restricted virulence factors may result in variation of the pathogenicity and environmental survival of different lineages [12, 50, 59].

Plasmid mediated acquisition of virulence genes was observed in *E. albertii*. The lineage restricted *hlyABCD* genes, the *iuc* gene cluster and the *lng* gene cluster were observed in MOB-suite predicted plasmids, indicating plasmid mediated acquisition, which was supported by previously studies in *E. coli* [46, 50, 59]. The two *E. albertii* isolates from bats harboured a plasmid with T2SS genes and the metalloprotease encoding *stcE* gene. T2SS genes are critical for the survival and pathogenicity of bacteria [60]. And *stcE* gene, which is located on pO157 plasmid, contributes to the intimate adherence of EHEC and atypical *S. boydii* 13 [61, 62]. Like plasmids, prophages were also found to have contributed to the acquisition of virulence genes in *E. albertii*. The non-LEE effector genes of the T3SS were observed in intact prophages, which were found to be significantly associated with G5 prophages defined in this study. A previous report that lambdoid prophages carried various T3SS secretion effectors supports this finding [11]. Altogether, plasmids and prophages play key roles in the transfer of virulence genes in *E. albertii* and may facilitate large changes in pathogenicity like those seen in the pathovars of *E. coli* [16].

### Plasmid mediated AR genes were significantly associated with STs and geographic regions

The predicted MDR rate in Chinese *E. albertii* isolates is astonishingly high (85.9%, 146/170), with 35.9% highly resistant isolates. These results are supported by previous phenotypic results, which found isolates resistant to up 14 clinically relevant drugs and 11 drug classes [14]. Importantly resistance was observed to clinically relevant drug classes including sulfamethoxazole-trimethoprim, cephalosporin, streptomycin and beta-lactam antibiotics [63]. There is an urgent need for surveillance and control of the spread of MDR and using MLST, we identified some STs that were associated with MDR *E. albertii* in China. ST4638, ST4479, ST4633 and ST4488 carried proportionally more MDR isolates and were mainly from China, which should facilitate the surveillance of the MDR. The MDR in North America and Europe is emerging and the MDR associated STs from these continents were different from those of China. This may be due to the different control strategies for antibiotic use in different countries. Plasmid transmission is the main pathway to acquire antibiotic resistance gene. In this study, we identified plasmid types that are significantly associated with MDR using both PlasmidFinder and MOB-Suite [23, 32]. The MDR associated plasmid types would facilitate the surveillance and control of MDR spread. Moreover, most of the L6 isolates harboured AR/MDR genes without predicted plasmids observed, which indicates potential new plasmids or prophages, or other means of MDR acquisition in L6.

### Conclusion

In this study, the population structure of *E. albertii* was elucidated based on 169 genomes from China and 383 genomes from other countries. There were 8 lineages identified, 7 of which (L2-L8) belonged to previously defined clade 2. Isolates from clinical infections were found in all lineages suggesting that much of *E. albertii* has some pathogenicity. However, the uneven distribution of many virulence factors suggests that the degree of pathogenicity may differ across the lineages. The predicted MDR rate and MDR gene profiles varied between regions, STs and CCs, with Chinese isolates and STs being predominantly MDR. Plasmid replicon and MOB types that were significantly associated with MDR were identified. *E. albertii* contained a large number of prophages and were divided into 5 groups, with G5 prophages found to have contributed to the acquisition of the T3SS non-LEE effector genes. Therefore, prophages and plasmids played key roles in creating the virulence and MDR repertoires of *E. albertii*. Our findings provided fundamental insights into the population structure, virulence variation and MDR of *E. albertii*.

## Abbreviations

EHEC: enterohemorrhagic *Escherichia coli*;
T3SS: type III secretion system;
LEE: enterocyte effacement;
Cdt: cytolethal distending toxin;
ETT2: type III secretion system 2;
Stx: Shiga toxin;
AR: antimicrobial resistance;
MDR: multidrug resistance;
NCBI: National Center for Biotechnology Information;
MLST: multi-locus sequence typing;
ST: sequence type;
CC: clonal complexes;
HPI: high pathogenicity island;
MVP: Microbe Versus Phage.

## Acknowledgements

Zigong CDC and L. L. acknowledge and commemorate Mr Yimao Miao for the guidance in all research studies in Zigong CDC, China. The authors thank Duncan Smith and Robin Heron from UNSW Research Technology Services for technical assistance.

## Authors’ contribution

L.L., M.P., R.L., Y.X. H.W. and Q.L. designed the study. H.W., L.Z., L.B., G.Y., Z.Z, Z.W., Y.X. and Q.L. collected the isolates and sequenced genomes. L.L., C.L. and H.Z. curated the data. L.L, R.L, M.P, C.L. and X.Z. analysed the results, R.L., M.P., Y.X. and L.B. provided critical analysis and discussions. L.L. wrote the first draft. M.P. and R.L revised the drafts. All authors approved the final manuscript.

