## Supplementary Figure S1-S5 for "Comparative genomics of Chinese and international isolates of *Escherichia albertii*: population structure and evolution of virulence and antimicrobial resistance"

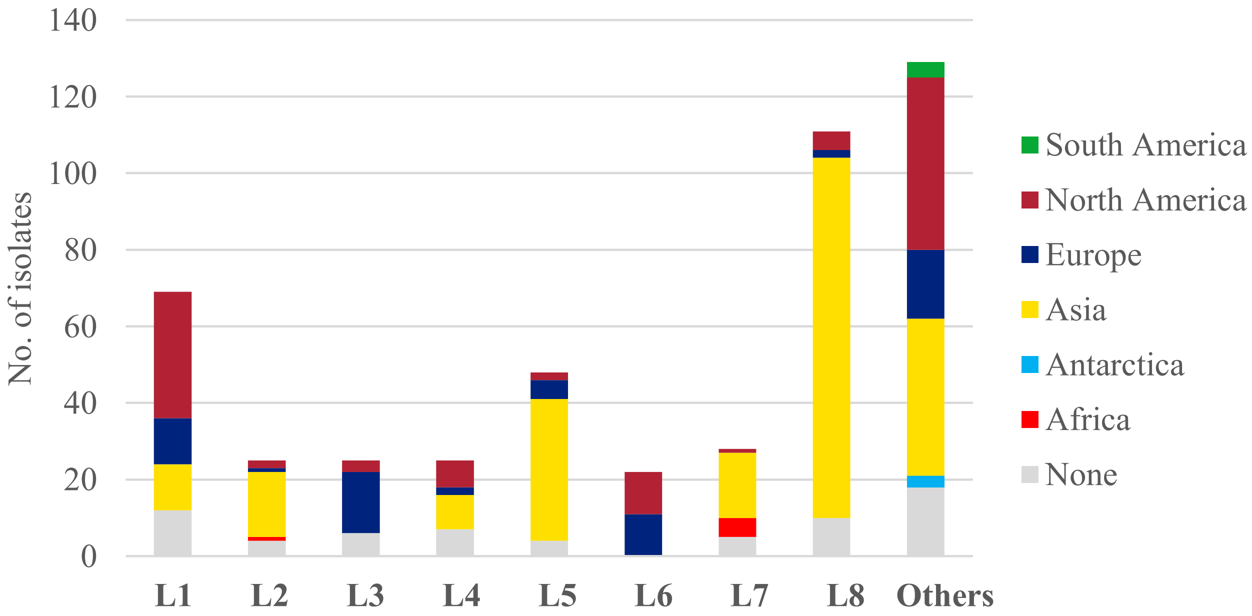


**Figure S1. Lineage distribution of *E. albertii* in different geographic regions.** Each column represented different lineages of *E. albertii* and the height the columns represented the number of isolates. Isolates from different geographic regions were shown in different colours. Asian *E. albertii* isolates were mainly observed in L2, L5, L7 and L8, and were not observed in L3 and L6. While the North American and Europe *E. albertii* isolates present in all of the lineages.

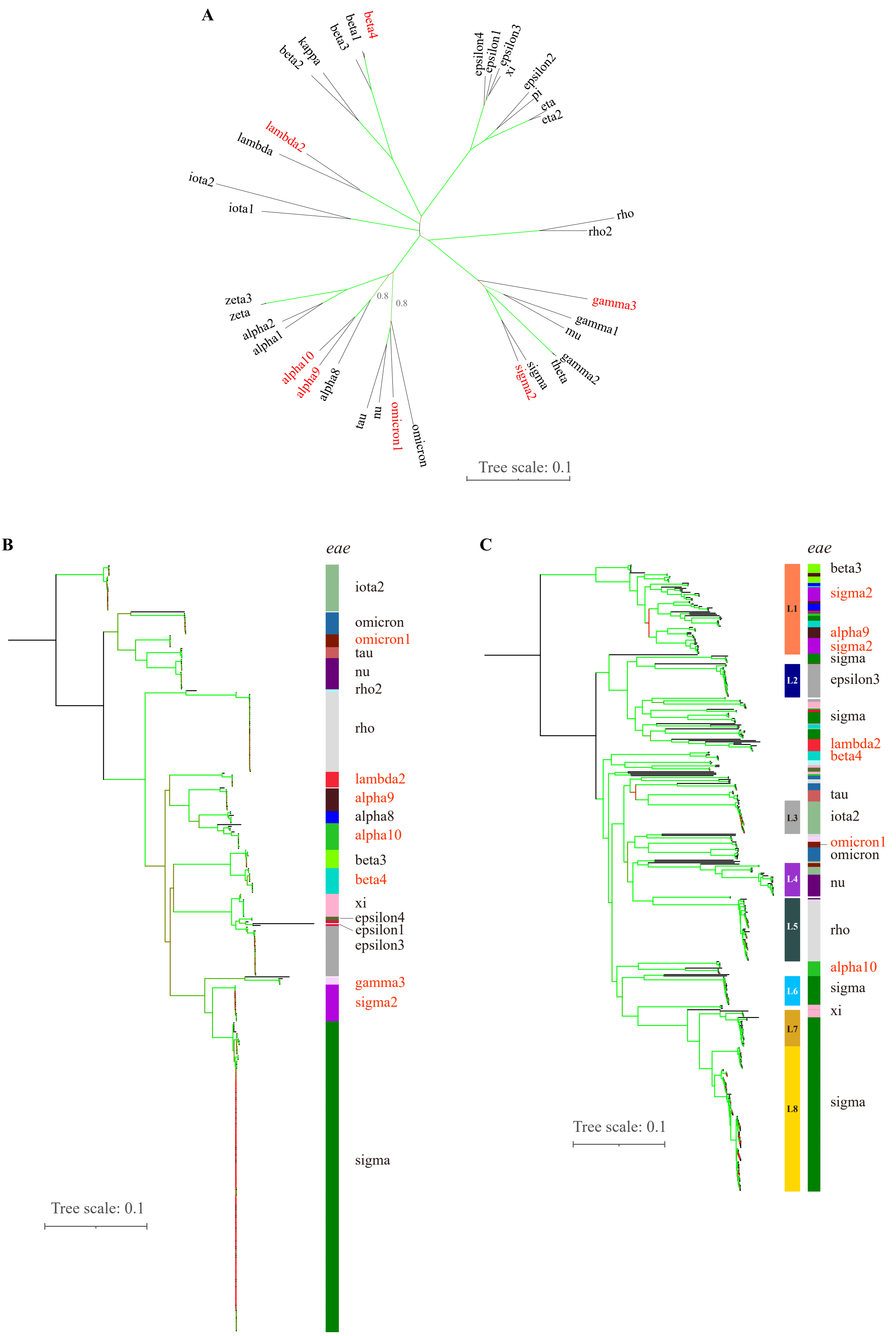
A
beta4
beta1
beta3
kappa
beta2
lambda2
lambda
iota2
iota1
epsilon4
epsilon1
epsilon3
xi
epsilon2
pi
eta
eta2
rho
rho2
zeta3
zeta
alpha2
alpha1
alpha10
alpha9
alpha8
0.8
0.8
tau
nu
omicron1
omicron
gamma3
gamma1
mu
gamma2
theta
sigma
sigma2
Tree scale: 0.1
B
eae
iota2
omicron
omicron1
tau
nu
rho2
rho
lambda2
alpha9
alpha8
alpha10
beta3
beta4
xi
epsilon4
epsilon1
epsilon3
gamma3
sigma2
sigma
Tree scale: 0.1
C
eae
L1
L2
L3
L4
L5
L6
L7
L8
beta3
sigma2
alpha9
sigma2
sigma
epsilon3
sigma
lambda2
beta4
tau
iota2
omicron1
omicron
nu
rho
alpha10
sigma
xi
sigma
Tree scale: 0.1

**Figure S2. Phylogeny of different *eae* gene subtypes in *E. albertii* and their distribution in different lineages.** (A) Neighbour joining (NJ) tree of 37 different subtypes of *eae* gene, including the 7 newly defined subtypes in red colour (1000 bootstrap, complete deletion). Branches with of 100% bootstrap support were highlighted in green colour. And branches with bootstrap of 80% support were labelled. (B) NJ tree of the *eae* gene from different *E. albertii* isolates with bootstraps of 1000. Each *eae* subtype was represented with a unique colour in the column strip. (C) Distribution of *eae* subtypes in different lineages of *E. albertii.* The phylogenetic tree was constructed based on the core SNPs of all 482 *E. albertii* isolates. The eight Fastbaps lineages were labelled in the strip. For the *eae* strip, different colours represented different *eae* subtypes corresponding to *eae* gene NJ tree in B.

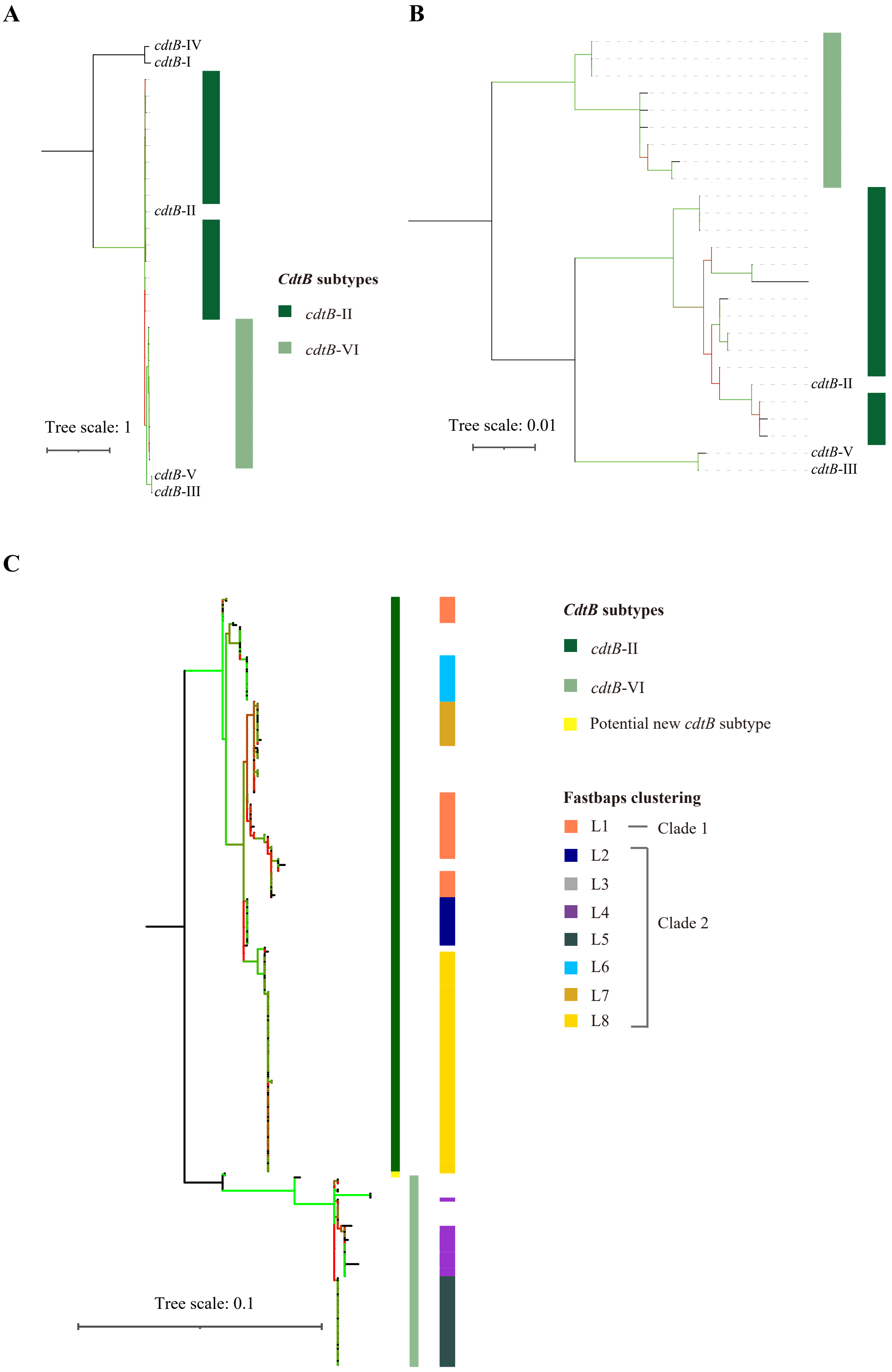

A
cdtB-IV
cdtB-I
cdtB-II
cdtB-V
cdtB-III
Tree scale: 1
CdtB subtypes
cdtB-II
cdtB-VI
B
cdtB-II
cdtB-V
cdtB-III
Tree scale: 0.01
C
CdtB subtypes
cdtB-II
cdtB-VI
Potential new cdtB subtype
Fastbaps clustering
L1
Clade 1
L2
L3
L4
L5
L6
L7
L8
Clade 2
Tree scale: 0.1

**Figure S3. Phylogeny of the *cdtB*-II and *cdtB*-VI from different *E. albertii* isolates.** (A) Maximum likelihood tree of different *cdtB* subtypes with 1000 bootstrap. The newly defined *cdtB*-VI subtype in *E. albertii* was phylogenetically closer to the *cdtB* II/III/V group. (B) Maximum likelihood tree of *cdtB* II/III/V/VI subtypes with 1000 bootstrap and complete deletion. This tree further supported *cdtB*-VI as a new subtype. (C) A maximum likelihood tree of the *cdtB*-II and *cdtB*-VI genes from 381 *E. albertii* isolates was constructed with bootstrap of 1000. The lineage distribution of the *cdtB* genes were highlighted in different colours. The *cdtB* gene from different lineages of *E. albertii* were clustered together, indicating dynamic evolution of *cdtB*-II and *cdtB*-VI along with the lineages of *E. albertii.*

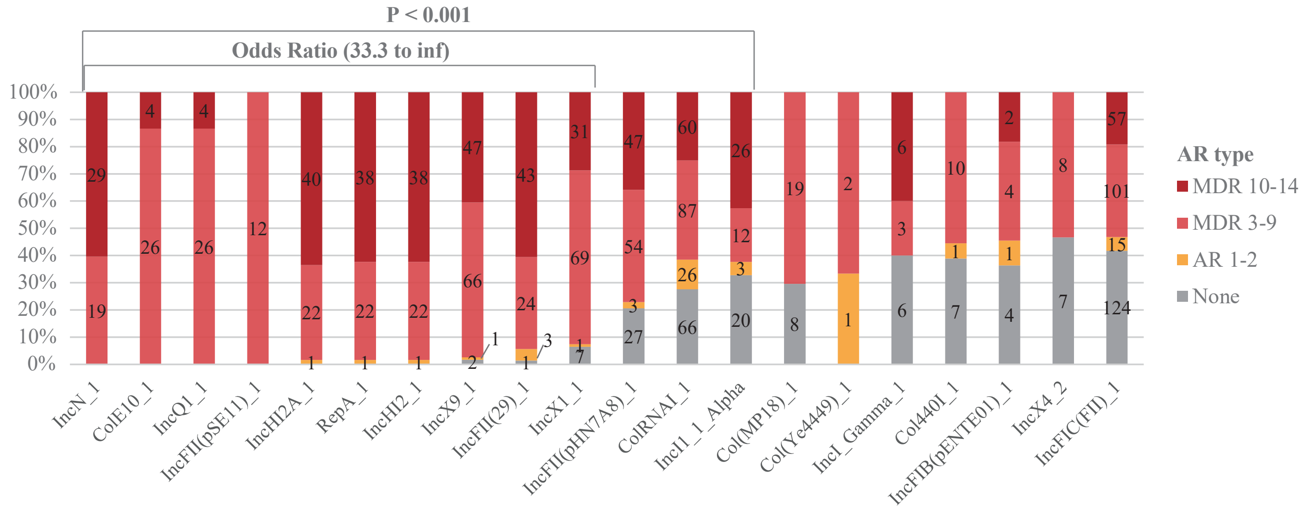


**Figure S4**. Plasmid subtypes that were associated with MDR. There were 13 replicon types significantly more likely to be observed in MDR isolates ($P < 0.001$). The first 10 replicon types were highly associated with MDR, the OR value of which were ranging from 38.2 to inf. However, the replicon specific genes may not be located on the same plasmid as MDR genes.

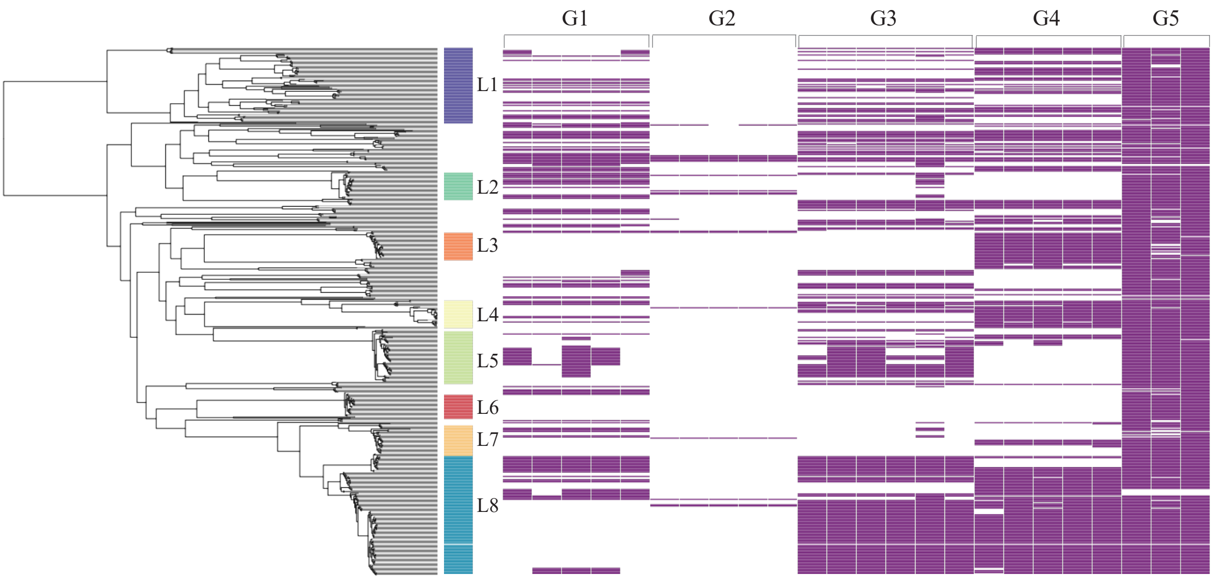


**Figure S5. Presence of prophage group specific genes in the assembly genomes of *E. albertii*.** For each group of prophages defined, three specific genes were identified with 100% sensitivity and specificity. The presence of a specific genes was marked in a coloured box. G5 prophage was predicted to be widely distributed in *E. albertii.* G2 prophages, which were mainly from the two bat source isolates, were predicted to be rarely present in *E. albertii*. Prophages belonged to G1, G3 and G4 were predicted to be absent in some lineages.

### Supplementary methods

#### The pangenome of *E. albertii* and phylogeny of the *eae* and *cdtB* genes.

High-quality assemblies were identified using the cutoffs of N50 <=290,000 base pairs, total length <=5.7 and >=4.3 million base pairs, and number of contigs <=450 contigs based on the output of Quast v5.0.2 [1]. A total of 422 high quality assemblies were included and were annotated using Prokka v1.12 [2]. The pangenome of *E. albertii* was defined using Roary v3.11.2 with the identity of 70% [3]. The *eae* and *cdtB* genes for each *E. albertii* isolate were identified from the output of Roary and Prokka. MEGA X was used to align the nucleotide sequences of *eae* and *cdtB* genes using muscle [4]. And neighbour joining trees were constructed with partial deletion (90%) and 1000 bootstrap repeats using MEGA X [4].

### Refernces


1. Gurevich A, Saveliev V, Vyahhi N, et al. QUAST: quality assessment tool for genome assemblies. Bioinformatics. 2013 Apr 15;29(8):1072-1075.
2. Seemann T. Prokka: rapid prokaryotic genome annotation. Bioinformatics. 2014 Jul 15;30(14):2068-2069.
3. Page AJ, Cummins CA, Hunt M, et al. Roary: rapid large-scale prokaryote pan genome analysis. Bioinformatics. 2015 Nov 15;31(22):3691-3693.
4. Kumar S, Stecher G, Li M, et al. MEGA X: Molecular Evolutionary Genetics Analysis across Computing Platforms. Mol Biol Evol. 2018 Jun 1;35(6):1547-1549.
